# Uncovering nick DNA binding by LIG1 at the single-molecule level

**DOI:** 10.1101/2024.03.28.587287

**Authors:** Surajit Chatterjee, Loïc Chaubet, Aafke van den Berg, Ann Mukhortava, Mitch Gulkis, Melike Çağlayan

## Abstract

DNA ligases repair the strand breaks are made continually and naturally throughout the genome, if left unrepaired and allowed to persist, they can lead to genome instability in the forms of lethal double-strand (ds) breaks, deletions, and duplications. DNA ligase 1 (LIG1) joins Okazaki fragments during the replication machinery and seals nicks at the end of most DNA repair pathways. Yet, how LIG1 recognizes its target substrate is entirely missing. Here, we uncover the dynamics of nick DNA binding by LIG1 at the single-molecule level. Our findings reveal that LIG1 binds to dsDNA both specifically and non-specifically and exhibits diffusive behavior to form a stable complex at the nick. Furthermore, by comparing with the LIG1 C-terminal protein, we demonstrate that the N-terminal non-catalytic region promotes binding enriched at nick sites and facilitates an efficient nick search process by promoting 1D diffusion along the DNA. Our findings provide a novel single-molecule insight into the nick binding by LIG1, which is critical to repair broken phosphodiester bonds in the DNA backbone to maintain genome integrity.

## Introduction

Genomic DNA is susceptible to damage from numerous sources, including endogenous and environmental factors (1,2). One potentially toxic form of DNA damage is the DNA strand breaks that occur naturally and continuously as intermediates during almost all cellular DNA transactions such as replication, repair, and recombination (3). If left unrepaired and allowed to persist, these strand breaks could result in potentially deleterious nicks in the DNA backbone, making them vulnerable to exonuclease-mediated degradation of DNA ends, leading to an increased frequency of recombination, and the formation of lethal double-strand breaks (4). The interruptions in the phosphodiester backbone of DNA are repaired by DNA ligases that convert nicks into phosphodiester bonds during discontinuous DNA synthesis on the lagging strand of the replication fork and at the final step of most DNA repair pathways (5). Therefore, the end joining by DNA ligase to generate an intact strain in almost every aspect of DNA transactions is critical for maintaining overall genome integrity (6,7).

All higher eukaryotic DNA ligases utilize ATP as a cofactor for joining two adjacent 5’-phosphate (P) and 3’-hydoxyl (OH) ends of nick in the energetically favorable consecutive three-step of the ligation reaction (3-6). During the first two steps, an adenosine 5’-monophosphate (AMP) moiety is covalently bound to an active site lysine residue of the ligase leading to the formation of LIG1-AMP intermediate (step 1), and then the AMP is transferred from the ligase to the 5’-PO_4_ end of a nick (step 2) resulting in the formation of the adenylated form of DNA (8,9). This DNA-AMP intermediate further activates the 5’-PO_4_ on the downstream strand for the nucleophilic attack by the upstream 3’-OH that displaces the adenylate moiety and catalyzes a phosphodiester bond formation between DNA ends during the final step 3 of the ligation reaction (10). Several enzymes that catalyze nucleotidyl transfers such as DNA ligases (ATP- and NAD^+^-dependent), RNA ligases, and RNA capping enzymes share this conserved ligation chemistry (11,12). Although the nick sealing reaction is highly conserved amongst the three kingdoms of life, and DNA ligation is the common and ultimate step of almost all critical DNA transactions in cells, how a DNA ligase recognizes and binds to its target nick substrate remains a critical knowledge gap in our understanding the molecular determinants of faithful DNA ligation.

The possible mechanism of nick binding has been suggested for DNA ligases from small viral DNA ligases, such as the Vaccinia and Chlorella viruses (13-19). DNA ligase adenylation at the active site lysine during the initial step of the ligation reaction is rapid and stable in the absence of nick DNA, which suggests that most ligase molecules in cells exist in this initial adenylated state (20). Furthermore, the requirement of a phosphate group at the 5’-end for proper nick sealing and weaker DNA nick binding by non-adenylated eukaryotic DNA ligase has been reported (15,16), suggesting that the initial formation of the LIG1-AMP intermediate is a prerequisite for nick recognition (21). The crystal structures of both NAD^+^- and ATP-dependent DNA ligases from various sources such as T4 DNA ligase, *Escherichia coli, Thermus thermophilus, Pyrococcus furiosus*, mycobacteria, and *Saccharomyces cerevisiae* have revealed a conserved mechanism that involves the distortion of the DNA helix upon nick binding which causes the region upstream of the nick to compress into an A-form helical conformation (22-29). Regarding the dynamics of nick sealing, the real-time fluorescence measurement of the ligation reaction using the TFAM reporter substrate with the T4 bacteriophage ligase demonstrated the first evidence of a multi-step DNA-binding and catalysis, suggesting a transition between two different conformational states of DNA ligase/nick substrate complex (23).

Human DNA ligases, DNA ligase I, IIIα, and IV, share a common catalytic core consisting of the oligonucleotide/oligosaccharide binding-fold (OB-fold) domain (OBD) and adenylation domain (AdD) that are well conserved in other DNA ligases and nucleotidyl transferases including RNA ligases and mRNA capping enzymes (4-6). The catalytic activity of the enzyme, both for ATP- and NAD^+^-dependent DNA ligases, is largely governed by the active site residues residing in the AdD and the OB-fold domains (20). Human DNA ligase 1 (LIG1), the main replicative ligase, carries out the nick sealing for Okazaki fragment maturation that occurs thousands to millions of times during each DNA replication cycle in eukaryotic cells (30). In addition, LIG1 joins the broken phosphodiester bonds to create an uninterrupted DNA strand by sealing a final nick product at the ultimate step after the excision of a damaged DNA base and subsequent re-synthesis of DNA during DNA repair pathways (31-33). Notably, the non-synonymous mutations (P529L, E566K, R641L, and R771W) in the *LIG1* gene have been described in the patients with LIG1 deficiency syndrome exhibiting immunodeficiency and cancer predisposition as well as aberrant DNA repair and replication (34,35).

The structure of LIG1 determined by X-ray crystallography revealed the N-terminal α-helical extension, which is recognized as the DNA binding domain (DBD) that stimulates nick sealing activity of the catalytic core (36). This first crystal structure of LIG1 also demonstrated that the AdD and OBD domains interact to form a continuous protein surface that engages the minor groove of DNA and this interaction with DNA alters the substrate conformation resulting in adoption of RNA-like A form helix, partially unwinding the DNA duplex and positions the DNA ends at the active site for end joining (36). Furthermore, the catalytic region adopts extended and asymmetric conformation in the absence of DNA and a large conformational change occurs between AdD and OBD domains that interact with DBD to form a ligase protein clamp-like architecture which encircles a DNA nick (36). According to the DNA binding model suggested based on the first structure of LIG1, the DBD domain binds non-specifically to double-stranded DNA, and then diffuses along the double helix to find a nick where the ligase adopts the ring-shaped active complex that encircles the 3’-OH and 5’-PO_4_ termini of the nick (36). Further structural studies revealed that LIG1 fidelity is mediated by Mg^2+^-dependent DNA binding (37). We also reported in our structures that LIG1 active site engages with nick containing mismatches distinctly depending on the architecture of 3’-primer/template base (38). Although the structure/function studies of LIG1 provided insights into the modular domain architecture of the ligase and how the ligase discriminates against unusual DNA ends, the dynamics of ligation reaction and the mechanism of the nick DNA binding by LIG1 have not been elucidated at the single-molecule level.

In addition to the common catalytic core, LIG1 contains a distinct N-terminal region including a proliferating cell nuclear antigen (PCNA) interacting peptide (PIP) box and a nuclear localization signal (NLS) that direct the ligase to participate in nuclear replication (39,40). This unstructured region also mediates LIG1 interactions with other repair and replication proteins such as PCNA trimer, clamp loader replication factor A (RFA) and C (RFC), Rad9-Rad1-Hus1 complex, a heterotrimeric clamp involved in cell cycle checkpoints as well as DNA polymerase (pol) β, a main base excision repair (BER) polymerase (41-44). Yet, the role of this non-catalytic region to nick DNA binding of LIG1 is entirely missing.

In the present study, we uncovered how LIG1 searches and binds to nick DNA at the single-molecule level. In addition, by comparing DNA binding modes of the full-length and C-terminal mutant of LIG1, we elucidated the importance of the N-terminal domain for binding to nicks on DNA. For this purpose, we used the C-Trap (LUMICKS) that combines optical tweezers, microfluidics, and a three-color confocal microscope to precisely visualize the positions of fluorescently labeled LIG1 on a biotinylated double-stranded (ds) lambda DNA substrate containing ten defined nick sites. Our findings, for the first time, revealed 1D diffusion mode of DNA binding and formation of long-lived DNA-LIG1 complex at the nick site. Furthermore, we uncovered the role of non-catalytic N-terminal region in facilitating 1D diffusion mode after the binding that leads to effective search and specific nick site binding. Our single-molecule characterization of LIG1/nick DNA dynamics provides a novel insight into the mechanism driving nick binding by a human DNA ligase, which is crucial stage of DNA ligation reaction that occur during DNA replication and repair.

## Methods

### Purification and fluorescent labeling of LIG1

Human DNA ligase 1 (LIG1) full-length (1-919 amino acids) and C-terminal (△261) proteins (pET-24b) with 6x his-tag were overexpressed in *E. coli* Rosetta (DE3) cells and grown in Terrific Broth (TB) media with kanamycin (50 μgml^−1^) and chloramphenicol (34 μgml^−1^) at 37°C as described (45-48). Once the OD_600_ reached 1.0, the cells were induced with 0.5 mM isopropyl β-D-thiogalactoside (IPTG) and overexpression was continued overnight at 20 °C. After centrifugation, the cells were lysed in the lysis buffer containing 50 mM Tris-HCl (pH 7.0), 500 mM NaCl, 20 mM imidazole, 2 mM β-mercaptoethanol, 5% glycerol, and 1 mM PMSF by sonication at 4 °C. The lysate was pelleted at 31,000 x g for 90 min at 4 °C. The cell lysis solution was clarified and then loaded onto HisTrap HP column in the binding buffer containing 50 mM Tris-HCl (pH 7.0), 500 mM NaCl, 20 mM imidazole, 2 mM β-mercaptoethanol, and 5% glycerol. The protein was then eluted with an increasing imidazole gradient (0-500 mM) at 4 °C. The collected fractions were subsequently loaded onto HiTrap Heparin in the binding buffer containing 20 mM Tris-HCl (pH 7.0), 50 mM NaCl, 2 mM β-mercaptoethanol, and 5% glycerol, and then eluted with a linear gradient of NaCl up to 1 M. LIG1 proteins were further purified by Superdex 200 10/300 column in the buffer containing 50 mM Tris-HCl (pH 7.0), 200 mM NaCl, 1 mM DTT, and 5% glycerol. Purity and concentrations of the resulting protein solutions were confirmed by SDS-PAGE gel analysis, and absorbance at 280 nm. LIG1 proteins were then fluorescently labeled using amine reactive NHS ester conjugated Alexa Fluor (AF)488 according to the manufacturer’s instructions (Thermo Fisher). The protein concentrations and labeling efficiency were finally determined by calculating the molar ratio of dye to protein within the solution by measuring absorbance at 280 nm and 488 nm.

### Confocal imaging in the C-Trap

Single-molecule experiments were performed on the C-Trap instrument combining three-color confocal fluorescence microscopy with dual-trap optical tweezers (LUMICKS) as described (49-53). The microfluidic flow-cell containing four distinct flow channels separated by laminar flow was moved by a computer-controlled stage to allow two optical traps to traverse the different laminar layers (Supplementary Figure 1). The single-molecule ligase/nick DNA binding experiments were performed in the buffer containing 1 mM HEPES (pH 7.4), 20 mM NaCl, 0.02% BSA (w/v), and 0.002% Tween 20 (v/v). The flow cell was first passivated with 0.05% (w/v) casein in PBS by flushing for 20 min and then incubating for an additional 10-20 min. The flow cell was then rinsed with the buffer for 20 min. For data collection, channels 1 and 2 were filled with 4.89 μm streptavidin-coated polystyrene beads and biotinylated lambda dsDNA that were diluted to 0.005% w/v and 0.05 ng/μl in the buffer, respectively. Channel 3 was used for trap calibration and channel 4 contained AF^488-^labeled LIG1 protein in the same buffer. Before starting the experiment, the protein channel was flushed for 10 min, incubated with LIG1 for another 30 min, and then rinsed with the buffer. While maintaining flow at a constant pressure of 0.3 bar, single beads were caught in both optical traps in channel 1 using a trap stiffness of 0.25 pN/nm. Then, the beads were moved to channel 2 for DNA capture. To tether DNA between the two traps, the bead in the left trap (upstream) was held in a constant position while the bead in the right trap (downstream) was moved closer and further away from the left bead. A characteristic increase in the force with increasing distance between the beads indicated the attachment of a DNA tether. A force-distance curve was measured for every tether and compared to the extensible wormlike chain model for dsDNA to verify that a single tether of dsDNA was caught. Once confirmed, the traps were moved to channel 4, and confocal line scanning of the nick DNA began. The Atto647N fluorophores on the DNA were excited intermittently to extend the lifetime of the fluorophores while AF^488^ was continuously excited to monitor the ligase/nicked DNA binding events.

#### Data Analysis

Data were analyzed with customer software from LUMICKS; tracking of binding events on kymographs was performed using Lakeview, and analysis was performed using Pylake. Confocal scans and force spectroscopy data were exported from the LUMICKS Bluelake acquisition software and processed using custom-written python scripts utilizing the Pylake package (54). Since the DNA tether is narrower than the pixel size of the confocal microscope, we employed a one-dimensional (1D) line scan which allowed faster frame rates than two-dimensional scanning. Binding density profiles were calculated by averaging over time the intensity of binding events of a kymograph. The coordinates of the nicks were computed by utilizing the known coordinates of the ATTO647N dyes. Profiles of multiple kymographs were combined by interpolating individual profiles and then adding the interpolated profiles together.

#### Diffusion coefficients

The first step for determining the diffusion coefficient was to track binding events. If a track displayed both diffusive and static behavior, it was split into two parts and the diffusion coefficient was determined for each part separately. Furthermore, we excluded events that were diffusing into the beads, or were hindered by other proteins, to ensure that only tracks from proteins diffusing freely are analyzed. The diffusion coefficient was computed for each tracked event using the Covariance Based Estimator, a fast and unbiased approach for determining diffusion coefficients (55).

#### Unbinding times

Unbinding times were calculated by tracking ligase binding events from start to end and extracting the duration of each tracked event. Binding durations from multiple kymographs were combined into one histogram. To determine the optimal model for the distribution of unbinding times, we fitted 1, 2, 3 and 4 exponentials using maximum likelihood fitting and computed the uncertainty of the fitted parameters using bootstrapping. We used the Bayesion information criterion (BIC) in combination with bootstrapping to determine which model was optimal (1, 2, 3 or 4 exponentials). Often, the model with the lowest BIC value is chosen as the optimal model. We noticed however that the bootstrapping distribution was not always unimodal for the model with the lowest BIC model, and that one of the components would be close to zero for some of the bootstrap distributions. Therefore, we selected the model that gives the lowest BIC value for which the bootstrapping distributions are still unimodal.

## Results

### Monitoring real-time LIG1 binding to nick DNA at the single-molecule level

For single-molecule characterization of the ligase/nick DNA binding, we used fluorophore-labeled LIG1 proteins full-length (1-919 amino acids) and the C-terminal region (261-919 amino acids) harboring the catalytic core composed of AdD, DBD, and OBD domains as well as the DBD domain (Supplementary Figure 1A). The biotinylated lambda double-stranded (ds) phage DNA construct (48.5 kbp) including ten predicted nick sites generated by Nt.BspQI nickase, two fluorophores (ATTO647N) at position 33,786 bp and 44,826 bp, was used and held between two 4.89 μm streptavidin-coated polystyrene bead (0.1% w/v) handles (Supplementary Figure 1B).

We performed experiments on the C-Trap which combines single-molecule fluorescent spectroscopy with an optical tweezer system and a multilaminar flow cell (Supplementary Figure 1C). During experiments, a single DNA tether is suspended between two polystyrene beads trapped by the optical tweezers to directly visualize the dynamics of nick DNA binding by AF^488-^labeled LIG1 (Supplementary Figure 1D). When a single DNA tether captured between two beads was confirmed, we moved the traps into the protein channel and began confocal scanning to observe real-time LIG1 binding to the DNA. Plotting the 1D confocal line scans as a function of time produces kymographs which show the position of AF^488-^labeled LIG1 molecules as a function of time along the nick DNA. We located the markers using a peak detection algorithm on the red channel and then used the known coordinates of the markers to identify the locations of the nicks. The construct was imaged using the integrated confocal system at 40 ms time resolution (Supplementary Figure 1E).

### LIG1 full-length exhibits multiple DNA binding behaviors that are enriched at nick sites

Individual kymographs demonstrated that LIG1 full-length protein binds both specifically and non-specifically to the nick dsDNA with a mix of both static and diffusive binding behaviors (Figure 1A). Interestingly, in several examples, a non-specifically bound LIG1 (Figure 1B, shown as orange arrow) initially either dissociates fast or exhibits 1D diffusive search on the DNA (Figure 1B, shown as pink arrow) followed by the formation of a static DNA-bound complex at a predicted nick site (Figure 1B, shown as yellow arrow). LIG1 can switch between 1D diffusive mode to a stable binding mode several times in the same trace, which demonstrates the free interconversion between these binding modes that LIG1 exhibits during nick searching (Figure 1B, shown as pink arrow). Several complete trajectories, displaying this initial remote binding to dsDNA, to facilitated diffusion towards a nick and finally to stable binding at the predicted nick site were observed. Analysis of the overall binding density along the DNA of individual kymographs further demonstrates that LIG1 is preferentially enriched at nick locations (Figure 1C).

**Figure 1.**
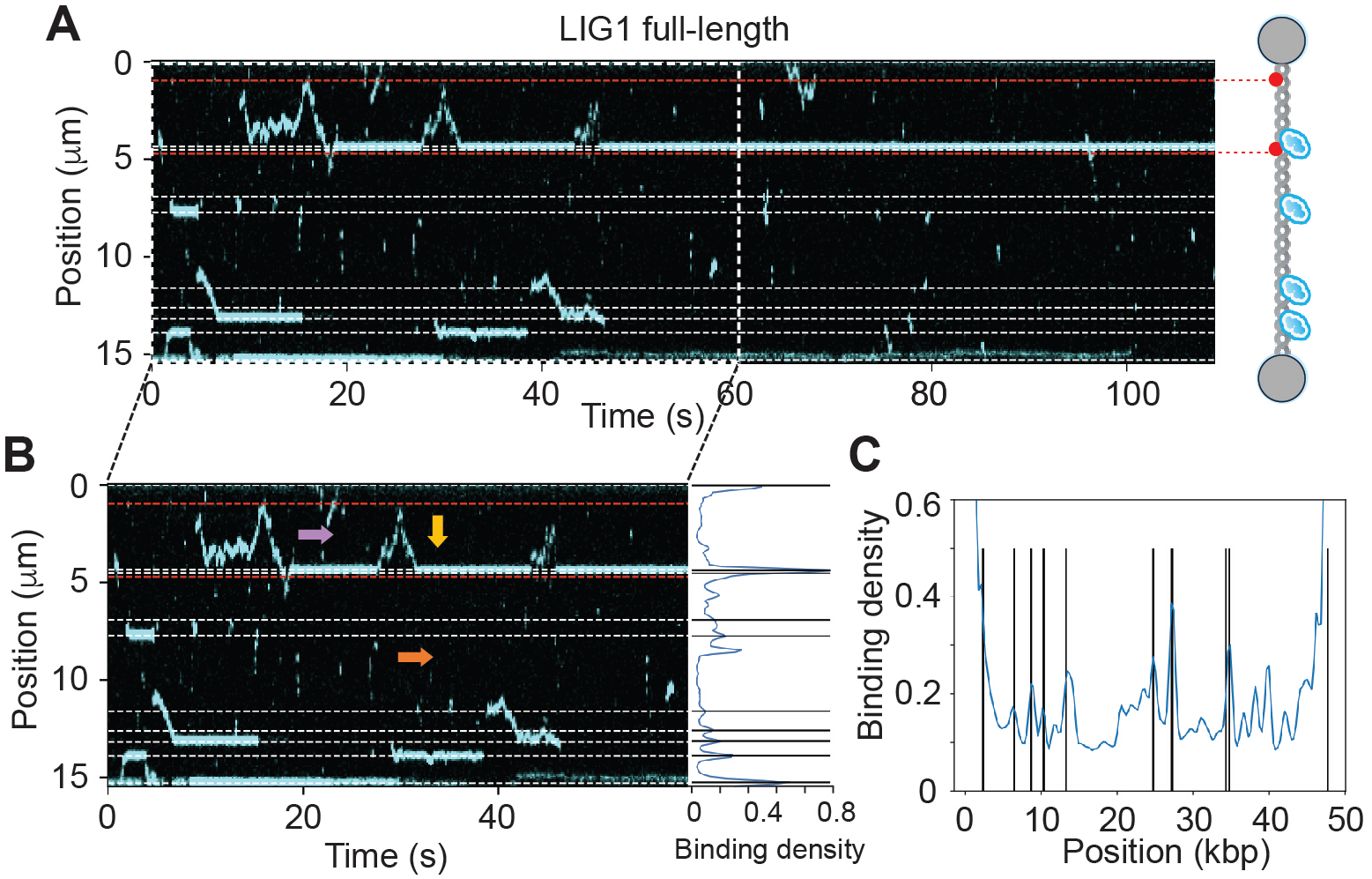
Single-molecule analysis reveals distinct modes of LIG1 binding to nick DNA. **(A)** A representative kymograph shows the binding positions of AF^488^-labeled LIG1 full-length protein (blue) on nick DNA as a function of time. Right panel; Schematic representation of the biotinylated lambda dsDNA with two marker fluorophores (ATTO647N) and bound proteins (blue). The DNA containing 10 predicted nick sites (dashed lines) is tethered between optically trapped beads. (**B**) Selected region from the kymograph in panel A indicates different binding modes with colored arrows. Non-specific DNA binding of LIG1 is shown as an orange arrow, while 1D diffusive binding on the DNA is depicted as a pink arrow, which is followed by the formation of a static DNA-bound complex at a predicted nick site shown as a yellow arrow. Right panel; Cumulative fluorescence binding intensity profile of the full kymograph (0-555 sec), where the peak intensities corresponding to bound LIG1 overlap well with the position of the predicted nick sites on the DNA (in kbp). **(C)** From the kymographs, the sum-over-time binding profile was computed by integrating all the signal over the entire length of the kymograph, then normalizing by the highest peak value. Profiles from three kymographs were combined for LIG1 full-length protein. Peaks correspond to the ligase-bound DNA and vertical lines represent the nick positions on dsDNA substrate (kbp).

Our results also demonstrated that LIG1 occasionally shows stable binding on a location that is not one of the predicted nicks (Supplementary Figure 2, shown as white arrow) probably due to the presence of an unpredicted nick site. We also observed that diffusive ligase cannot pass static ligase (Supplementary Figure 2, shown as blue arrow) and that the LIG1 was also capable of 1D diffusion past several predicted nick sites (Supplementary Figure 2, shown as green arrow).

### LIG1 C-terminal binding to nick DNA is non-specific

To elucidate the role of non-catalytic N-terminal region for binding of LIG1 to nick DNA, we next used the LIG1 protein harboring C-terminal region (Supplementary Figure 1A) and observed robust binding activity throughout the DNA devoid of specific binding on the nick locations (Figure 2). LIG1 C-terminal protein displays a single binding mode throughout the DNA, regardless of predicted nick sites. Indeed, the individual kymographs showed non-preferential binding throughout the DNA and when bound at or near the nicks, contrary to the full-length LIG1 (Figure 1). Occasionally, the LIG1 C-terminal protein seems to diffuse over a nick without switching to a static binding mode at the nick (Figure 2A, lower panel, shown as pink arrow). Overall binding density along the DNA showed little to no preferential enrichment of LIG1 C-terminal protein at nick sites (Figure 2B), suggesting that the N-terminal non-catalytic region plays a significant role for promoting specific binding to target nick sites by LIG1.

**Figure 2.**
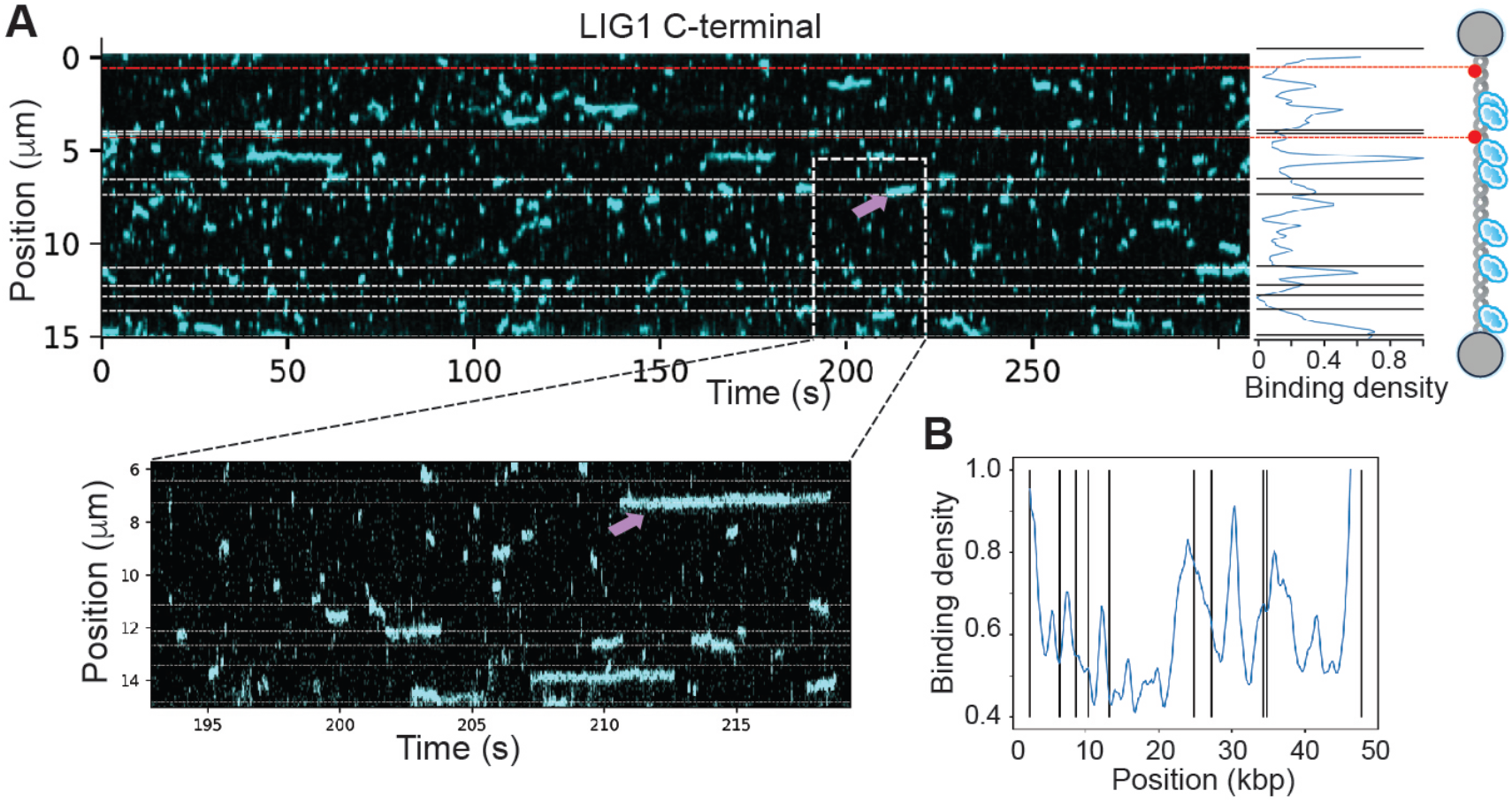
LIG1 C-terminal mutant binds non-specifically to nick DNA. **(A)** The kymograph (top panel) displays single molecule dynamics AF^488^-labeled LIG1 C-terminal (blue) lacking N-terminal region of the protein binding to nick DNA. Bottom panel shows selected region of the kymograph displaying multiple non-specific off-target binding and few binding events at predicted nick sites (dashed lines). **(B)** Combined binding profiles from five kymographs for the LIG1 C-terminal protein binding exhibits no binding enrichment near the nick sites.

### Role of non-catalytic N-terminal region for nick binding by LIG1

To further characterize the role of the N-terminal domain of LIG1 for the binding to nick DNA, we estimated the binding lifetimes of DNA-bound LIG1 full-length and C-terminal proteins (Figure 3). The binding lifetime was defined as the lifetime of the full event, with diffusive and static binding modes combined. The binding kinetics was calculated combining the binding lifetimes of the full-length protein from six kymographs. The data can best be fit with three exponential time scales (Figure 3A and Supplementary Figure 3). For the LIG1 C-terminal protein, the binding kinetics was calculated combining binding lifetimes from three kymographs. We predominantly observed fast transient binding and the lifetimes can best be fit with two exponential time scales (Figure 3B and Supplementary Figure 4). For the LIG1 full-length protein (Figure 3C), we observed a frequency of 220 binding events in 10 min time scale with binding lifetimes of 0.08 sec (58%), 0.74 sec (27%), and 11 sec (15%). This suggests that the majority of LIG1 dissociates fast, while 15% of the ligase stays bound for very long about 11 sec. For the LIG1 C-terminal protein (Figure 3C), we found an average frequency of 122 binding events per min with binding lifetimes of 0.25 sec (78%) and 2 sec (22%).

**Figure 3.**
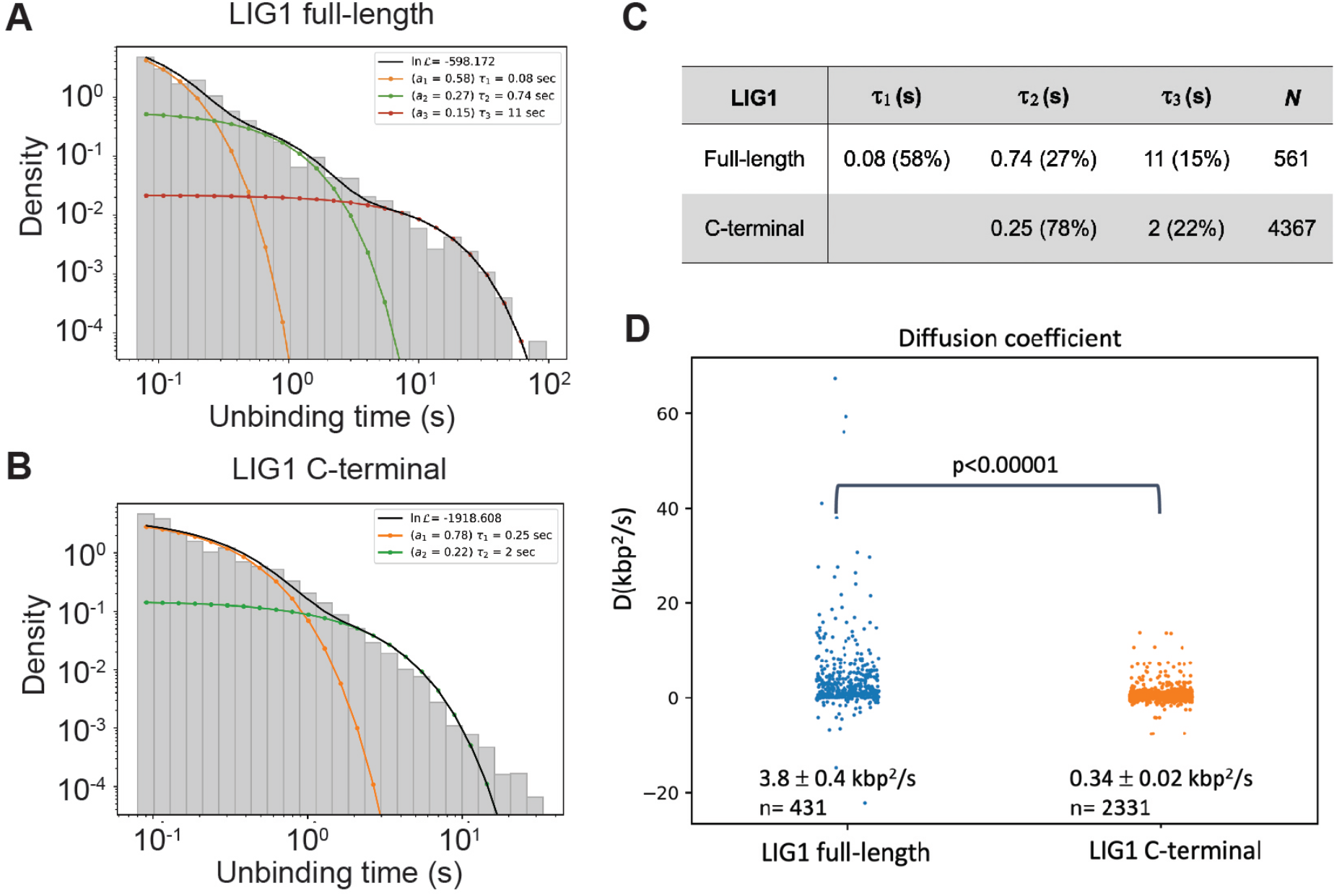
Comparisons for the unbinding times of LIG1 full-length and C-terminal proteins. **(A-B)** The unbinding times were fitted with three exponentials for LIG1 full-length (A) and two exponentials for C-terminal (B) timescales. **(C)** Binding lifetimes extracted by fitting 2 or 3 exponential functions to the distribution of binding lifetimes for LIG1 full-length and C-terminal proteins. *N* refers to a number of total binding events. **(D)** Distribution of the diffusion coefficients of LIG1 full-length (blue circles) and C-terminal (orange circles) proteins, estimated diffusion coefficients are determined as mean +1 SEM. The distributions were compared using a Welch’s T test.

Furthermore, we calculated the diffusion coefficient for the 431 diffusive events of LIG1 full-length and the 2331 diffusive events of LIG1 C-terminal observed during our experiment (Figure 3D). We found that the rate of diffusion for LIG1 full-length (3.8 ± 0.4 kbp^2^/s) was ∼10-fold higher than that of LIG1 C-terminal (0.34 ± 0.02 kbp^2^/s), suggesting that the N-terminal domain has an important role in mediating fast and efficient diffusion of LIG1 on nick DNA. Together, overall results suggest that, without the N-terminal domain, the LIG1 C-terminal could bind non-specifically and remains bound to the DNA for significantly shorter periods compared to the full-length protein.

## Discussion

Visualizing DNA-binding proteins while interacting with their substrates at the single-molecule level elucidates individual dynamics and binding modes of the DNA-binding proteins with their substrates which are averaged out in bulk assays (56). Single-molecule studies have been extensively applied to examine in real-time how DNA repair proteins detect specific lesions and bind to their targets at extraordinary detail using purified proteins with a fluorescence tag and living cells or cell extracts (53,56-65). LIG1 is the fundamental enzyme to maintain the structural integrity of the genome as it functions in almost all DNA transactions such as DNA replication, repair, and recombination that all generate strand breaks in the phosphate backbone of DNA (6). As such DNA intermediates can threaten the loss of genetic information and could cause the introduction of deleterious chromosomal mutations if left unrepaired, it’s important to understand the details of diverse interactions that LIG1 employs to detect and bind to nick sites on genomic DNA.

The nick sealing activity of LIG1 is attributed to the catalytic region consisting of AdD and OBD domains that harbor the critical active site residues for catalysis (20). LIG1 protein contains an extended protease sensitive and unstructured N-terminal region that directs the ligase to be recruited to nuclear DNA replication foci via nuclear localization signal (30). Also, the PIP box motif containing eight amino acids mediates its interaction with PCNA and is also important for Okazaki fragment synthesis (32). LIG1 interacts with many other replication and repair proteins through this distinct N-terminal region (32). Furthermore, the serine/threonine residues residing within this region undergo posttranslational modifications and are phosphorylated during cell cycle progression by cyclin-dependent kinase and casein kinase II. It has been shown in mammalian cell models *in vivo* that N-terminal deletion mutant of LIG1 cannot rescue lethal LIG1-null phenotype and is essential for cellular function and viability to DNA damage inducing agents (33). Although this N-terminal region is dispensable for catalytic activity of LIG1 *in vitro*, its role in nick sensing and binding is largely unknown.

Our results demonstrate that after initially binding to DNA, the LIG1 full-length either dissociates fast or exhibits 1D diffusion to search a nick site. Whereas the LIG1 C-terminal protein binds non-specifically throughout the DNA and exhibits extremely diminished ability to diffuse along the DNA and find a nick site. Our findings suggest the role of LIG1 N-terminal region in promoting 1D diffusion along the DNA. Furthermore, a significant percentage of the DNA-bound full-length LIG1 (15%) exhibited long-lived residence time on the DNA and LIG1 binding is enriched at nick sites, suggesting the formation of stable DNA-LIG1 complex at the nick sites. For the LIG1 C-terminal protein, the binding lifetimes are significantly shorter without any enrichment at the nick sites, suggesting non-specific and comparatively less stable LIG1 C-terminal/DNA interaction. Previous single-molecule studies also reported different modes of DNA interactions and collaborative dynamic process during the initial DNA damage search by repair proteins. For example, nucleotide excision repair protein UvrA exhibits 3D diffusion in solution and remains bound on DNA, upon association with UvrB, a significant fraction of UvrA becomes motile and exhibits 1D dissociation (65). Similarly, the binding time and diffusion mechanism of DNA glycosylases varies significantly during the damage search process and that, an increased binding lifetime suggests that the protein is bound to a damage site (57,58). Poly [ADP-ribose] polymerase 1 (PARP1) has been also reported to exhibit mostly 3D diffusion to identify its substrate and the dissociation of DNA-bound-PARP1 is facilitated in the presence of AP-Endonuclease 1 (APE1) that induces 1D diffusion to enable downstream repair processes (61).

Based on our results, we propose a working model for nick DNA binding mode of LIG1 (Figure 4). LIG1 binds to the dsDNA both specifically and non-specifically and can diffuse to the nick site to form a stable complex at the nick site. Majority of the initial LIG1 full-length binding occurs through fast transient interaction, facilitating efficient nick target search process. Whereas only a significant percentage (15%) of DNA-bound LIG1 forms stably bound complex on the DNA. LIG1 without N-terminal region, however, upon initial binding non-specifically to DNA, cannot diffuse along the DNA and remains bound for significantly shorter time before eventually dissociating from the DNA. Altogether, our data demonstrate the efficient mechanism of LIG1 binding to nick targets and how the N-terminal region helps to search nick targets by promoting 1D diffusion along the DNA as well as by minimizing unnecessary LIG1 binding to undamaged regions.

**Figure 4.**
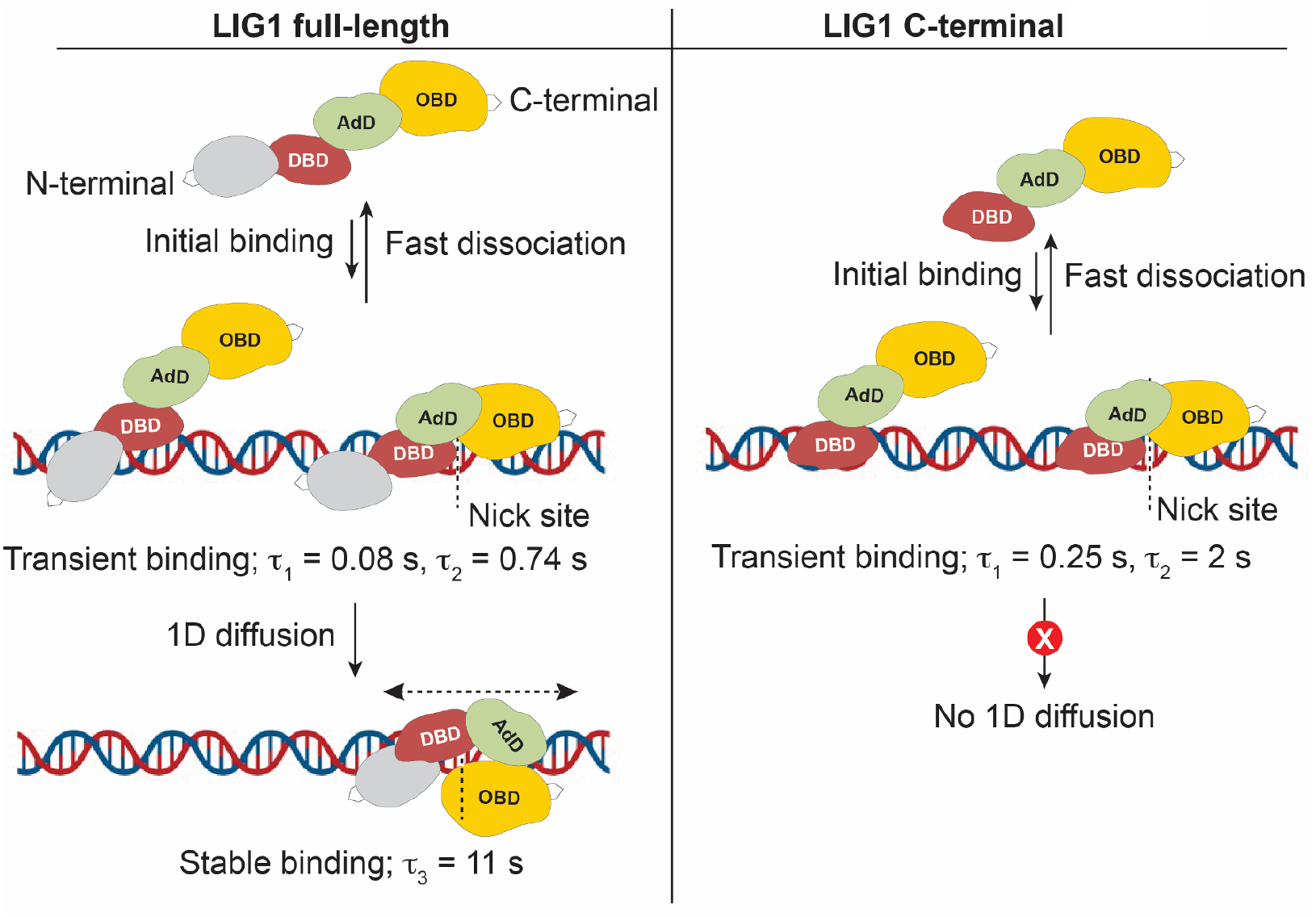
Proposed model of LIG1 nick DNA binding. Initial transient binding of LIG1 full-length protein occurs from solution and is enriched at the nick site. LIG1 can then diffuse 1D to search the nick site and form a stable complex. Without the N-terminal region, the LIG1 C-terminal binds non-specifically throughout the DNA only transiently without the ability to diffuse.

Our study represents a novel insight into the mechanism of nick recognition and binding by LIG1 at the single-molecule level, which is critical to better understand how strand-breaks are repaired at the final ligation step of almost all DNA repair pathways and during the maturation of Okazaki fragments in DNA replication. These insights into nick DNA binding mechanism may potentially lead to the development of effective and specific DNA ligase inhibitors for targeted therapeutic applications (66).

