## Supplementary Figures for "Uncovering nick DNA binding by LIG1 at the single-molecule level"

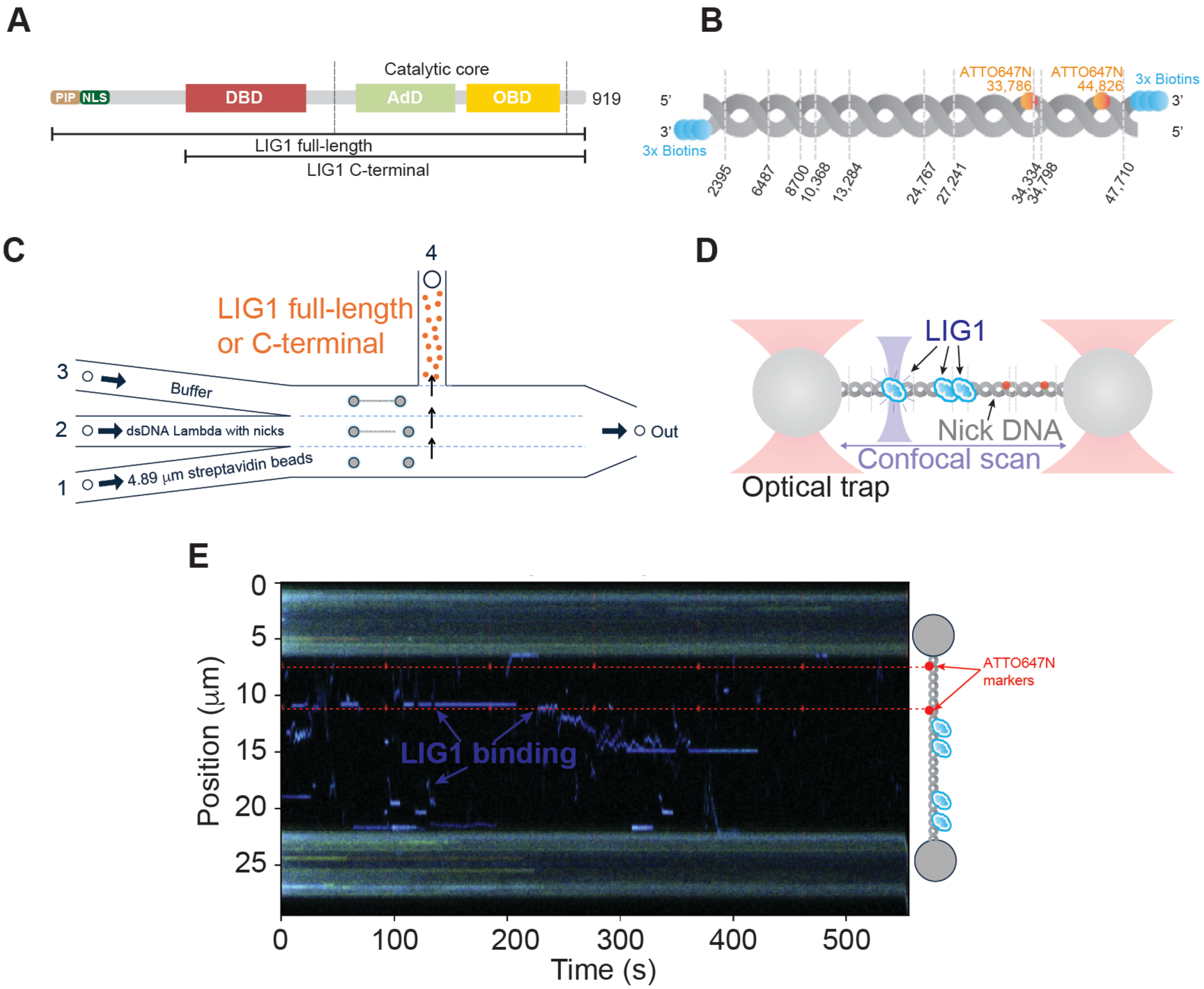
A
PIP
NLS
DBD
Catalytic core
AdD
OBD
919
LIG1 full-length
LIG1 C-terminal
B
ATTO647N 33,786
ATTO647N 44,826
3x Biotins
5'
3'
2395
6487
8700
10,368
13,284
24,767
27,241
34,334
34,798
47,710
C
4
LIG1 full-length or C-terminal
3
Buffer
2
dsDNA Lambda with nicks
1
4.89 μm streptavidin beads
Out
D
LIG1
Nick DNA
Confocal scan
Optical trap
E
Position (μm)
Time (s)
LIG1 binding
ATTO647N markers
0
5
10
15
20
25
100
200
300
400
500

**Supplementary Figure 1. (A)** The protein domain organization of LIG1 for the full-length (1-919 amino acids) and C-terminal region (261-918). The DNA-binding domain (DBD, red) and the catalytic core composed of the adenylation (AdD, green) and Oligonucleotide-binding or OB-fold (yellow) domains and N-terminal region including the nuclear localization signal (NLS) and proliferating cell nuclear antigen interacting peptide (PIP) box are indicated. **(B)** Schematic representation of the 48,502 bp biotinylated l-dsDNA with ten nicks used in this study. The two fluorophores (ATTO647N) at position 33,786 bp and 44,826 bp were used to identify the locations of the nicks. **(C)** Schematic describes the workflow of the C-Trap instrument combining three-color confocal fluorescence microscopy with dual-trap optical tweezers (LUMICKS). A microfluidic flow-cell containing four distinct flow channels separated by laminar flow was moved by a computer-controlled stage to allow two optical traps to traverse the different laminar layers. Channels 1 and 2 are filled with 4.89 μm streptavidin-coated polystyrene beads and biotinylated lambda dsDNA, respectively. Channel 3 is used for trap calibration in the reaction buffer and channel 4 contains $AF^{488-}$labeled LIG1 (full-length or C-terminal). When a single DNA tether was confirmed, the traps were moved into the protein channel 4 where confocal scanning of LIG1 on DNA was measured. **(D)** Schematic shows the DNA captured between two optically trapped polystyrene beads for confocal scanning of $AF^{488-}$labeled LIG1 binding on nicked DNA. **(E)** Example kymograph displaying the ATTO647N markers and $AF^{488}$-labeled LIG1 full-length protein (blue) binding events on the DNA as a function of time. The markers (ATTO647N) are first located using a peak detection algorithm on the red channel and then the known coordinates of the markers were used to identify the locations of the nicks.

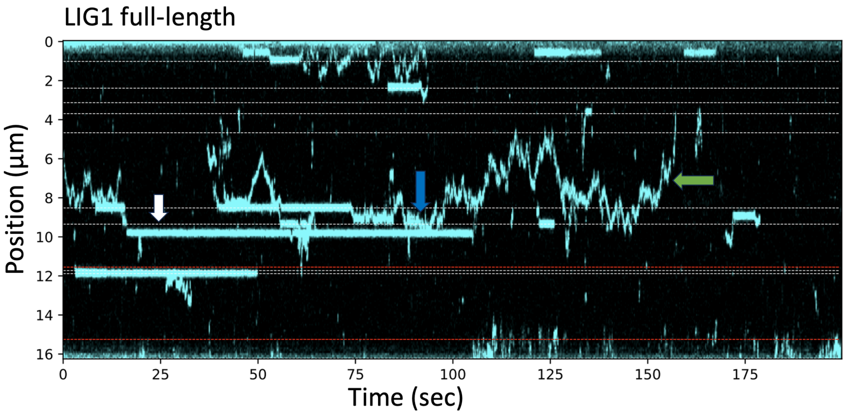


**Supplementary Figure 2.** Example of kymograph shows the single molecule dynamics of AF$^{488}$-labeled LIG1 full-length protein (blue) binding on nick DNA as a function of time. LIG1 occasionally shows stable binding on a location that is not one of the predicted nicks (white arrow). The diffusive ligase cannot pass static ligase (blue arrow) and is capable of 1D diffusion past several predicted nick sites (green arrow).

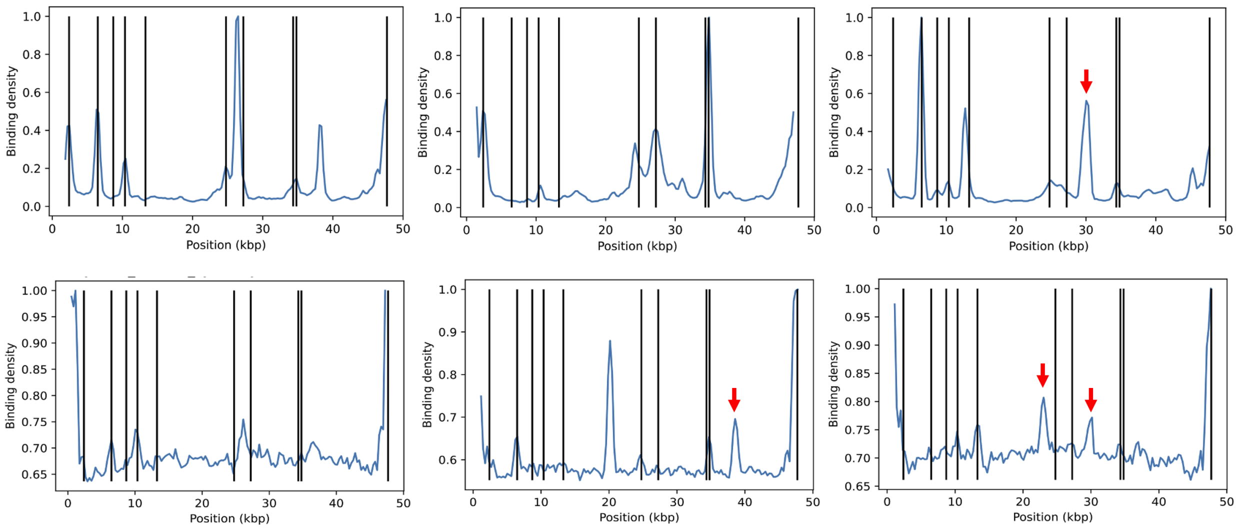


**Supplementary Figure 3. Nick DNA binding kinetics of LIG1 full-length.** DNA binding profiles for individual kymographs for 50 sec duration/each of LIG1 full-length. Peaks (blue) correspond to the ligase bound to DNA and the vertical lines (black) represent nick coordinates and their positions on dsDNA substrate (kbp). Red arrows indicate non-specific and off-target DNA bindings by LIG1. These off-target binding sites could correspond to binding to unpredicted nick sites. Overlaps of peaks with vertical lines correspond to LIG1 bound to one of the ten predicted nick sites on dsDNA.

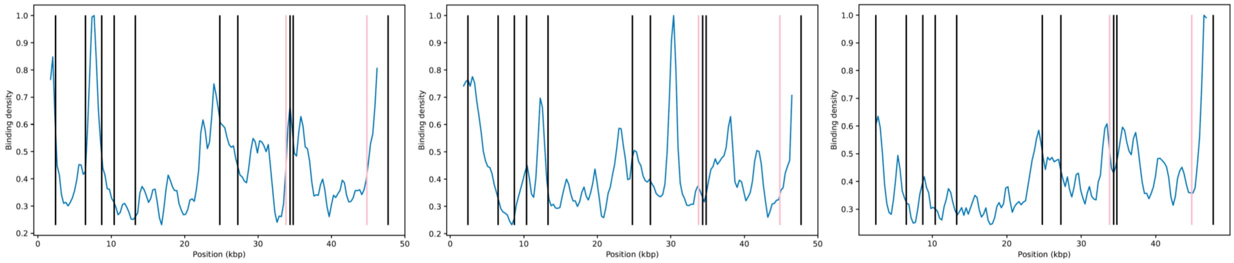


**Supplementary Figure 4. Nick DNA binding kinetics of LIG1 C-terminal protein.** DNA binding profiles for individual kymographs for 50 sec duration/each of LIG1 C-terminal protein. Peaks (blue) correspond to the ligase bound to DNA and the vertical lines (black) represent nick coordinates and their positions on dsDNA substrate (kbp).
